# SOX2 can associate with chromatin directly by binding to DNA or indirectly via association with other chromatin-bound proteins

**DOI:** 10.64898/2026.09.02.748901

**Authors:** Briana D. Ormsbee Golden, Kabhilan Mohan, Ethan P Metz, Phillip J Wilder, Jimmie Persinger, Soumi Basu, Daisy V. Gonzalez, Tahir H. Tahirov, Donald W. Coulter, Sandipan Brahma, Angie Rizzino

## Abstract

It is widely assumed that SOX2 regulates gene expression and facilitates the opening of chromatin by binding directly at SOX motifs. To test this assumption, we created a SOX2 DNA binding mutant to determine whether other regions of SOX2 contribute to gene target specificity. When exogenously expressed in cells, this SOX2 mutant [SOX2(G76P)], like elevated unmodified SOX2, dramatically alters the transcriptome, but it does so by regulating vastly different gene sets and gene networks than SOX2. Consistent with their differential effects on the transcriptome, ChIP-seq analysis demonstrates that SOX2 and SOX2(G76P) associate primarily with different genomic loci, and motif analysis indicates that SOX2 binds primarily at SOX motifs, whereas SOX2(G76P) associates with chromatin at non-SOX motifs, including AP-1 motifs. Additionally, ATAC-seq analysis indicates that SOX2 substantially increases chromatin accessibility, but SOX2(G76P) does not. The findings presented lead to the conclusion that SOX2(G76P) associates with chromatin indirectly by a “piggyback” mechanism through its association with other chromatin-associated proteins, including AP-1 complexes. Remarkably, we also show that SOX2 and SOX2(G76P) each associate with a subset of the same gene loci that contain several different DNA motifs, including AP-1 motifs, but no high confidence SOX motifs. Overall, our findings provide new perspectives on SOX2 and lead to two important conclusions: 1) selection of gene targets by SOX2 is not solely determined by its DNA binding domain, and 2) SOX2 not only associates with chromatin directly by binding to SOX motifs but can also associate with a subset of gene loci indirectly through its association with other chromatin-associated proteins.

## Introduction

It is generally accepted that most transcription factors determine when, where, and how strongly genes are expressed by binding with high specificity to relatively short consensus DNA sequences (DNA motifs) located in gene regulatory regions. A defining feature of transcription factors is a highly structured DNA-binding domain (DBD). In the case of SOX transcription factors, each member contains a highly structured high mobility group (HMG) box that is homologous to the HMG box of the founding member of the family, SRY (sex-determining region Y gene) (Gubbay et al. 1990; Sinclair et al. 1990; Rizzino and Wuebben 2016). In addition to highly structured DBDs, transcription factors possess intrinsically disordered regions (IDRs) (Wright and Dyson 1999), including their transactivation domains, that interact with a diverse array of transcriptional machinery, such as the chromatin remodeling complex SWI/SNF (Jonas et al. 2025), to regulate gene transcription. In the case of the stem cell transcription factor SOX2, a member of the SRY box (SOX) transcription factor family (Gubbay et al. 1990; Sinclair et al. 1990), nearly the entire protein except for its DBD is predicted to be disordered by AlphaFold (Jumper et al. 2021). This includes its ∼40 amino acid N-terminus and its ∼200 amino acid C-terminus, which contains its transactivation domain (Nowling et al. 2000). The importance of IDRs is highlighted by studies linking several diseases to mutations in IDRs that are believed to disrupt essential protein-protein interactions (Wong et al. 2020). Significantly, recent studies have found that the gene specificity of some transcription factors *in vivo* is heavily influenced by their IDRs that mediate protein-protein interactions with other transcriptional machinery (Brodsky et al. 2020; Jonas et al. 2025; Abidi et al. 2026). Surprisingly, for a few transcription factors, their IDRs are sufficient to drive target specificity (Brodsky et al. 2020; Abidi et al. 2026). The extent to which other regions of additional transcription factors contribute to target specificity is currently unknown, and this includes SOX2.

SOX2 is expressed in over 20 different types of human cancer (Wuebben and Rizzino 2017). Importantly, the levels of SOX2 in these cancers varies widely. In neuroendocrine cancers, such as neuroendocrine prostate cancer and ASCL1 subtype small cell lung cancer, SOX2 is highly expressed (Tenjin et al. 2020; Bernal et al. 2024). Moreover, SOX2 levels have been shown to rise significantly during tumor progression due, at least in part, to mutations in tumor suppressor genes, such as RB1 and TP53 (Mu et al. 2017; Ku et al. 2017). SOX2 levels have also been shown to vary within a given tumor and have been found to be elevated in their cancer stem cell populations (Wuebben and Rizzino 2017; Metz and Rizzino 2019). This is also true in animal models, such as the GEM model of sonic hedgehog (SHH) medulloblastoma (MB), where SOX2 was detected in less than 5% of the tumor population (Vanner et al. 2014). Significantly, this minor SOX2 high cell population, which proliferates infrequently, can reconstitute the tumor when chemotherapy is halted. Interestingly, SOX2-positive cancers appear to have hijacked an important normal function of SOX2 related to growth control. During mammalian development, high levels of SOX2 are expressed in the infrequently proliferating fetal stem cell populations of multiple developing tissues where high levels of SOX2 have been linked directly to the infrequent proliferative status of their fetal stem cell populations (Hagey and Muhr 2014; Hagey et al. 2018).

To help understand how elevated levels of SOX2 modulate tumor cell proliferation, we engineered more than a dozen tumor cell lines, representing seven different human tumor cell types, to elevate SOX2 from a doxycycline-inducible promoter (Cox et al. 2012; Wuebben et al. 2016; Metz et al. 2020a, 2020b). Without exception, elevation of SOX2 in each of the tumor cell lines leads to a substantial reduction in cell proliferation *in vitro*. When three of these tumor types (prostate, pancreatic ductal adenocarcinoma, and MB) were examined *in vivo,* elevation of SOX2 induced a reversible state of tumor growth arrest in each case (Metz et al. 2020b).

Significantly, when doxcycline was removed, SOX2 levels quickly returned to basal levels and tumor growth resumed at rates comparable to their untreated control counterparts. More recently, we determined that increasing SOX2 in tumor cells leads to substantial remodeling of the transcriptome with over six thousand genes up- and downregulated at the RNA level (Metz et al. 2022; Ormsbee Golden et al. 2024). Given the impact on the transcriptome when SOX2 levels are altered, it is vital to understand the full scope of how SOX2 directs its specificity.

In this study, we created a SOX2 DNA-binding mutant, referred to as SOX2(G76P), to examine whether regions of SOX2 that are not directly involved in DNA binding also contribute to target specificity. SOX2(G76P) was generated by replacing glycine with proline at residue 76 in human SOX2. This substitution disrupts a critical α-helix in its DBD (Reményi et al. 2003). Using cells engineered for inducible expression of SOX2(G76P), we set out to answer four questions regarding SOX2(G76P): 1) Is it biologically active? 2) Is it able to remodel the transcriptome? 3) Does it associate with chromatin? 4) Does it regulate the same set of genes as SOX2? During these studies, we also examined whether SOX2 and SOX2(G76P) can associate with chromatin at some of the same gene loci that lack SOX motifs.

## Results

### Association of mutant SOX2 with DNA *in vitro* and its effects on cell proliferation

To gain a broader understanding of the extent to which SOX2 contributes to cell proliferation and the regulation of gene transcription through alternate modes of target specificity, we created a subtle SOX2 DNA binding mutant. Based on the crystal structure of mouse SOX2 (mSOX2), its DBD consists of three α-helices, second of which (amino acids 72 to 81) binds within the minor groove of DNA (Reményi et al. 2003). Human SOX2 is two amino acids shorter than mSOX2 in its N-terminal region, and it is notable that the 79 amino acid sequence of the DBDs of mSOX2 and human SOX2 are identical. To disrupt the DBD, we replaced the glycine in mSOX2 at residue 78 with proline to create mSOX2(G78P) and replaced the glycine in human SOX2 at residue 76 with proline to create SOX2(G76P) (Supplemental Fig. S1A). Replacing a glycine with a proline is expected to disrupt the second α-helix of the DBD by introducing a “kink” in the helix. To determine how this change altered SOX2, we initially compared the ability of mSOX2 and mSOX2(G78P) to bind to DNA *in vitro*. For this purpose, Flag-tagged mSOX2 and Flag-tagged mSOX2(G78P) were transiently expressed in 293T cells, and their cell extracts were used to test their *in vitro* binding to DNA. This was done by performing an electrophoretic mobility shift assay with a DNA probe containing two SOX motifs found in the HDAC2 gene (Boyer et al. 2005). mSOX2 formed two prominent SOX2:DNA complexes (Supplemental Fig. S1B). The faster and slower migrating complexes are consistent with DNA probes bound by one and two mSOX2 molecules, respectively. Each complex shifted to slower migrating complexes in the presence of the M2 flag antibody (Supplemental Fig. S1B). In contrast, mSOX2(G78P) did not appear to form a specific DNA protein complex. Instead, it formed a much slower migrating faint band that did not shift to a slower migrating band in the presence of the M2 flag antibody, and this band matches what we observed with the empty vector. These results indicate that the ability of mSOX2(G78P) to bind to DNA is impaired significantly.

To determine how this mutant affects the ability of human SOX2 to function at the cellular and genomic level, we engineered the human medulloblastoma cell line, ONS76, for Doxycycline (Dox)-inducible expression of SOX2(G76P) to create i-SOX2(G76P)-ONS76 cells as described in the Methods section. ONS76 cells were selected for this study because we had already extensively characterized the effects of elevating SOX2 on the transcriptome and the growth of SOX2-inducible ONS76 cells (i-SOX2-ONS76). In this regard, we previously demonstrated that inducible elevation of SOX2 in i-SOX2-ONS76 cells leads to growth inhibition (∼40%) (Metz et al. 2020b). Here we demonstrate that elevating SOX2(G76P) in i-SOX2(G76P)-ONS76 cells also induces a dose-dependent decrease in cell proliferation (Fig. 1A, B).

**Figure 1.**
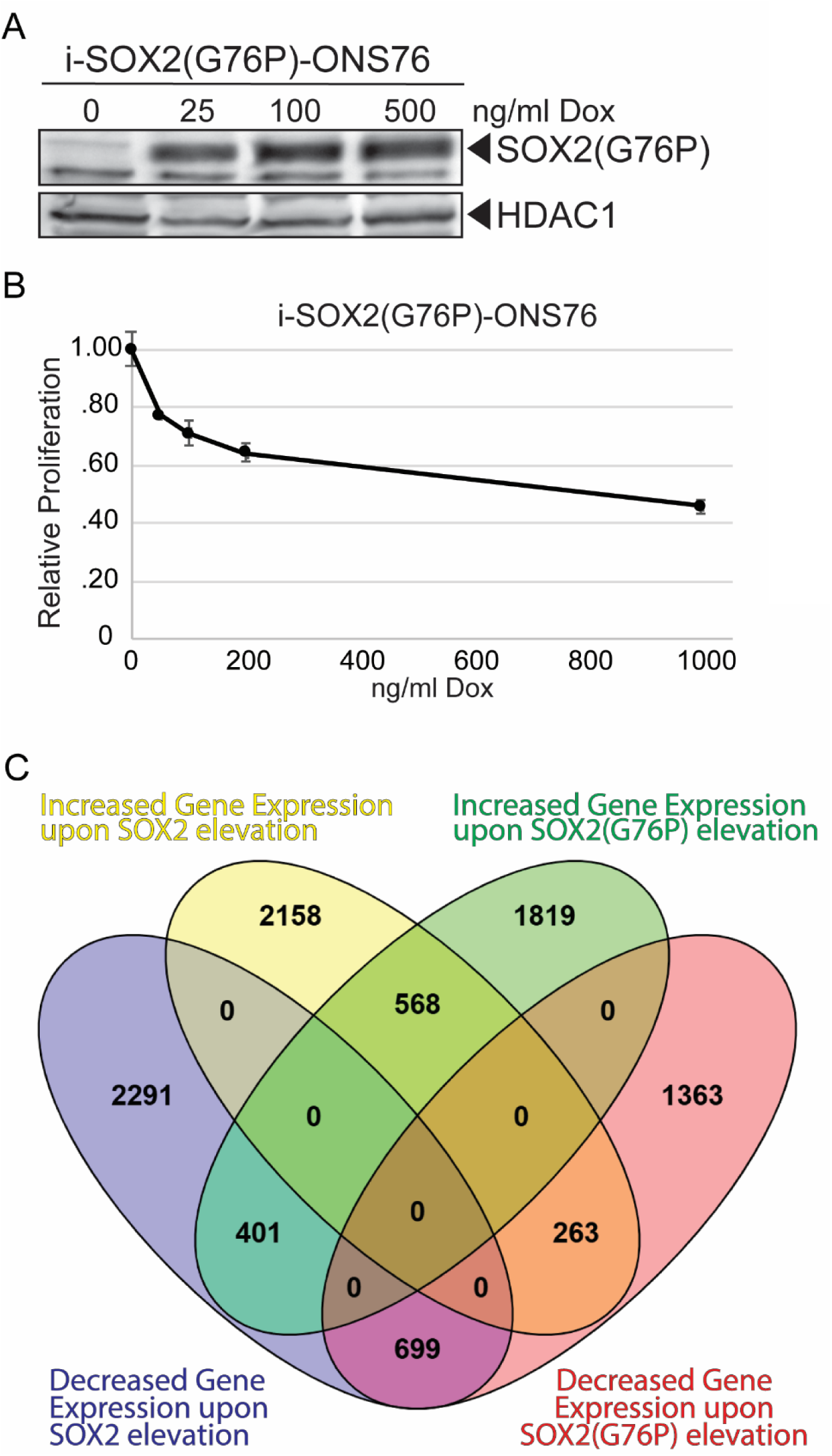
Characterization of i-SOX2(G76P)-ONS76 cells upon SOX2(G76P) elevation. (A)i-SOX2(G76P)-ONS76 control cells and cells induced to elevate SOX2(G76P) using increasing doses of Dox for 48 hours were processed for whole cell protein isolation and western blot analysis. HDAC1 served as a loading control. (B) MTT assay conducted to determine changes in proliferation after exposure to increasing concentrations of Dox for 96 hours. (C) Venn diagram of genes differentially expressed (q-value < 0.05; Log2 of -\+ 0.5) between control and Dox-induced i-SOX2-ONS76 and i-SOX2(G76P)-ONS76 cells.

### Differential Effects of SOX2 and SOX2(G76P) on the transcriptome

To extend the comparison between SOX2 and SOX2(G76P), we compared their effects on the transcriptome. Previously, we demonstrated that elevating SOX2 in i-SOX2-ONS76 cells upregulated over 2,900 genes and downregulated over 3,300 genes at the RNA level (Metz et al. 2022). Like the effect of elevating SOX2, elevating SOX2(G76P) significantly altered the transcriptome – over 2,700 genes were upregulated and over 2,300 genes were downregulated (Fig. 1C). However, only 568 genes were upregulated by both SOX2 and SOX2(G76P), and only 699 genes were downregulated by both SOX2 and SOX2(G76P) (Fig. 1C). Furthermore, Gene Ontology analysis demonstrated that SOX2 and SOX2(G76P) regulated vastly different biological processes (Supplemental Fig. S2A). Additionally, the transcription factors/machinery that target the genes exhibiting increased and decreased expression in response to elevation of SOX2 or SOX2(G76P) were significantly different (Supplemental Fig. S2B). Genes exhibiting increased expression when SOX2 was elevated in i-SOX2-ONS76 cells were identified as target genes in previous ChIP-seq experiments for chromatin remodeling factors: MBD3, SMARCA4, and p300 (Supplemental Fig. S2B). As noted above, the genes that increased in response to SOX2(G76P) expression were largely different than those increased by SOX2, including genes identified previously as targets of CJUN and ATF3 (Supplemental Fig. S2B), which can functionally partner to affect gene expression (Liu et al. 2016). Together, these findings demonstrate that both SOX2 and SOX2(G76P) significantly alter the transcriptome, but they do so by regulating vastly different sets of genes and gene networks.

### Association of SOX2 and SOX2(G76P) genome-wide

To examine whether SOX2 and SOX2(G76P) behave differently at the genomic level, we initially compared their ability to associate with chromatin. For this purpose, i-SOX2-ONS76 and i-SOX2(G76P)-ONS76 cells were cultured in the absence or presence of Dox before chromatin was isolated. Western blot analysis demonstrated that substantial amounts of both SOX2 and SOX2(G76P) could be extracted from chromatin prepared from Dox-induced i-SOX2-ONS76 and i-SOX2(G76P)-ONS76 cells, respectively (Supplemental Fig. S3A). Next, we examined whether SOX2 and SOX2(G76P) associate with similar genomic loci by performing ChIP-seq with a SOX2 antibody. Two replicates of each condition were generated using i-SOX2-ONS76 and i-SOX2(G76P)-ONS76 cells cultured without or with Dox. For both the control (without Dox) i-SOX2-ONS76 and control i-SOX2(G76P)-ONS76 cells, we observed ∼2,000 peaks (Fig. 2A). Given the large increase in SOX2 and SOX2(G76P) association with chromatin, it was not surprising that there was over a 4-fold increase in the number of peaks when SOX2(G76P) was elevated in i-SOX2(G76P)-ONS76 cells and over a 6-fold increase in the number of peaks when SOX2 was elevated in i-SOX2-ONS76 cells (Fig. 2A). Additionally, for both SOX2 and SOX2(G76P) most peaks were observed in the gene body, intron, and distal intergenic regions (Supplemental Fig. S3B). Importantly, motif analysis of individual samples demonstrated that although SOX2 ChIP-seq analyses detected association of SOX2 and SOX2(G76P) at some of the same motifs, they associate primarily with different DNA motifs (Supplemental Fig. S3C). As expected, SOX motifs were the most common motifs observed in both control i-SOX2-ONS76 and control i-SOX2(G76P)-ONS76 cells where endogenous SOX2 is expressed (Supplemental Fig. S3C). Similarly, SOX motifs were the most common motifs identified in the peaks obtained from the Dox-induced i-SOX2-ONS76 cells. In strong contrast, this was not the case for SOX2(G76P) peaks from the Dox-induced i-SOX2(G76P)-ONS76 cells. Importantly, FRA1, ELK1, ELK4, CTCF, and BORIS motifs were observed at high significance in the elevated SOX2(G76P) peaks, whereas SOX motifs were not observed (Supplemental Fig. S3C). These results indicate that SOX2 and SOX2(G76P) associate predominately with different genomic sequences.

**Figure 2.**
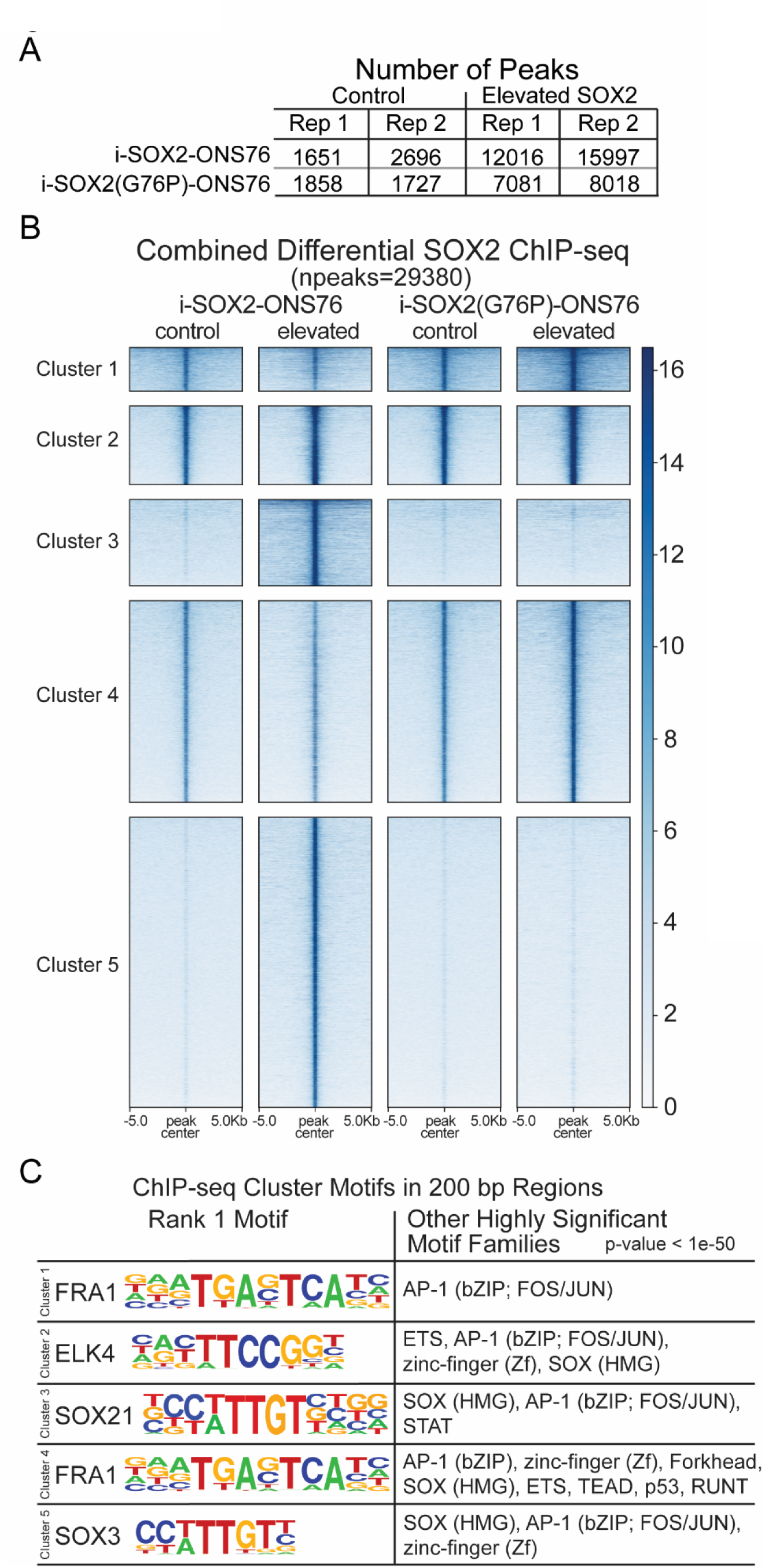
SOX2 and SOX2(G76P) ChIP-seq Analysis. (A) Changes in the number of genomic peaks 48 hours after SOX2 and SOX2(G76P) were elevated as determined by ChIP-Seq with a SOX2 antibody using two biological replicates for both conditions (control or Dox-induced elevation of SOX2 – either wild-type or the G76P mutant form) in each cell line. (B) Heatmap of unsupervised clustering of genomic regions from a combined differential between control and Dox-induced (elevated) i-SOX2-ONS76 and i-SOX2(G76P)-ONS76 ChIP-seq profiles with control i-SOX2-ONS76 serving as the reference. This analysis was performed for both biological replicates, which were highly similar. Only a single replicate for each condition is displayed. (C) The top motif as well as other highly significant (p-value < 1e-50) motif families identified using HOMER in 200 bp regions within each of the ChIP-seq clusters.

Significant differences between SOX2 and SOX2(G76P) are also evident when unsupervised clustering is used to compare genomic regions bound by SOX2 and SOX2(G76P) that change when their respective cells are DOX-induced (Fig. 2B). As expected, the clustering of genomic regions was similar for control i-SOX2-ONS76 cells and control i-SOX2(G76P)-ONS76 cells. Importantly, elevation of SOX2 in Dox-induced i-SOX2-ONS76 cells modestly increased association with genomic regions in cluster 2 and dramatically increased association with genomic regions in clusters 3 and 5. For clusters 3 and 5, SOX motifs were the top ranked motifs (Fig 2C). In strong contrast, expression of SOX2(G76P) in Dox-induced i-SOX2(G76P)-ONS76 cells led to little or no increase in association with genomic regions in clusters 3 and 5, whereas there was increased association with genomic regions in clusters 1, 2 and 4, in which the top ranked motifs were FRA1, ELK4 and FRA1 motifs, respectively (Fig. 2C). Collectively, these findings demonstrate that SOX2 and SOX2(G76P) associate predominantly with different sets of genomic regions and associate with different DNA motifs.

### Differential effects of SOX2 and SOX2(G76P) on chromatin accessibility

To probe for additional differences between SOX2 and SOX2(G76P), we performed ATAC-seq, because SOX2 is known to increase chromatin accessibility (Soufi et al. 2015). For these studies, two replicates for each condition were generated. Consistent with the known pioneering activity of SOX2, elevation of SOX2 in Dox-induced i-SOX2-ONS76 cells led to a large increase in the number of chromatin-accessible peaks genome wide (Fig. 3A). Specifically, increasing the level of SOX2 raised the basal number of ∼177,500 chromatin accessible peaks by nearly 60,000, for an increase of ∼33%. In strong contrast, expression of SOX2(G76P) led to only a small increase of ∼4,500 chromatin accessible peaks, for an increase of ∼2.5%. This large difference between SOX2 and SOX2(G76P) is consistent with previous studies showing that chromatin accessibility induced by transcription factors like SOX2 is initiated by their binding directly to nucleosomal DNA (Soufi et al. 2015; Dodonova et al. 2020). Further analysis of the ATAC-seq data by unsupervised clustering of genomic regions indicates that elevating SOX2 increases chromatin accessibility of some genomic regions (Fig. 3B clusters 3-5), while decreasing the accessibility of others (Fig. 3B clusters 1 and 2). In contrast, unsupervised clustering revealed very limited changes in chromatin accessibility patterns upon elevating SOX2(G76P). To extend the analysis, we assessed the chromatin accessibility of each of the SOX2 and SOX2(G76P) ChIP-seq genomic clusters shown in figure 2B (Supplemental Fig. S4). As expected, there were no demonstrable increases in chromatin accessibility within the ChIP-seq clusters that exhibited increased association of SOX2(G76P) when it was expressed in the Dox-induced i-SOX2(G76P)-ONS76 cells (Supplemental Fig. S4 clusters 1-3). In contrast, the ChIP-seq clusters that exhibited substantial increases in SOX2 binding when SOX2 was elevated in i-SOX2-ONS76 cells corresponded with increases in chromatin accessibility (Supplemental Fig. S4 clusters 4 and 5). Together, these studies indicate that increases in SOX2, but not SOX2(G76P), lead to substantial increases in chromatin accessibility.

**Figure 3.**
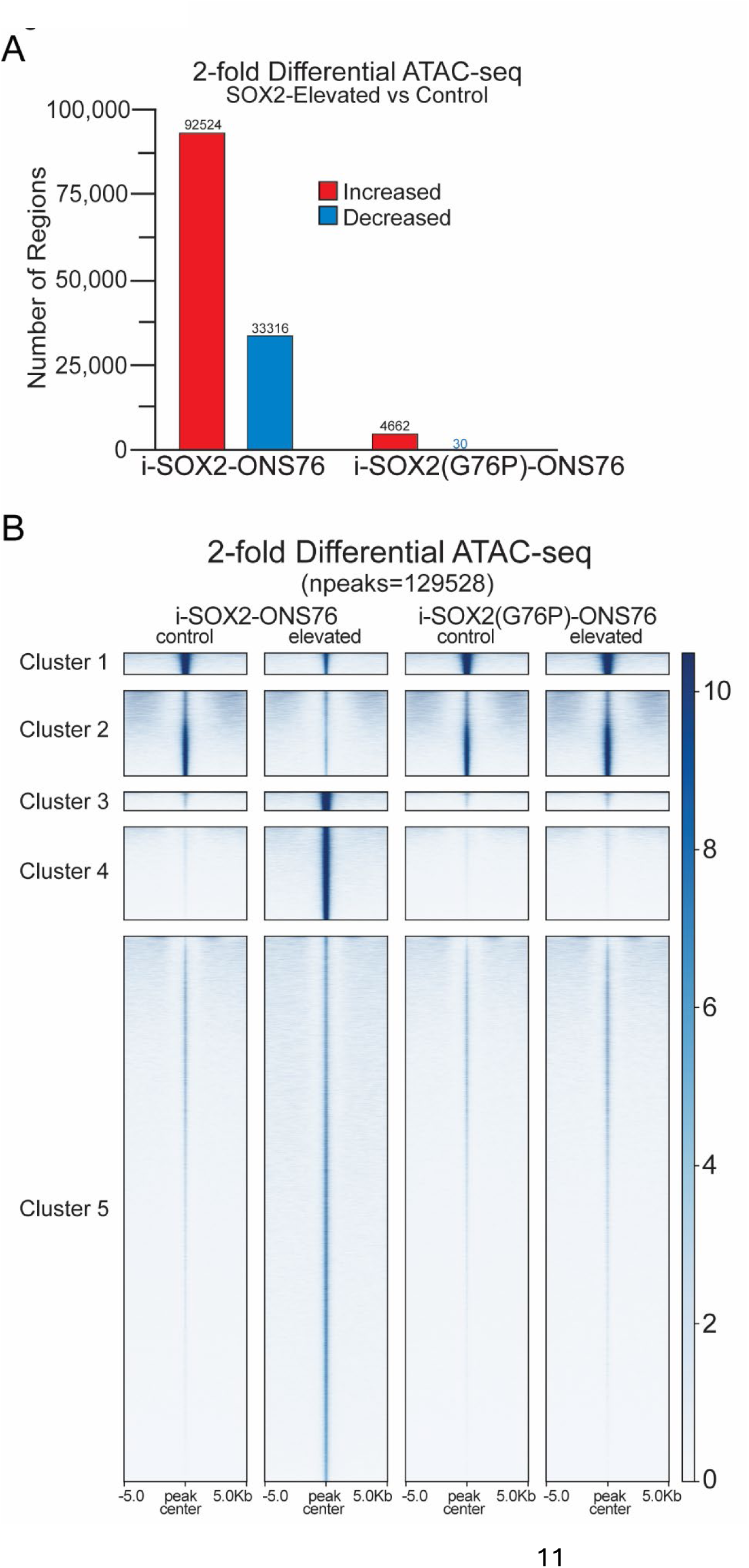
Changes in chromatin accessibility when SOX2 or SOX2(G76P) were elevated. (A) The number of genomic regions in the human genome (hg38) that exhibited a 2-fold or more increase (red) or decrease (blue) of high-throughput sequencing, representing the change in the amount of transposase-accessible chromatin when SOX2 or SOX2(G76P) was elevated compared to their control counterparts for the i-SOX2-ONS76 or i-SOX2(G76P)-ONS76 cell lines, respectively.(A)Heatmap of clustered genomic regions that exhibited >2-fold differential signals utilizing ATAC-seq results from control i-SOX2-ONS76 cells as the reference for comparison to ATAC-seq of Dox-treated i-SOX2-ONS76 cells after 48 hours, and control or Dox-treated i-SOX2(G76P)-ONS76 cells after 48 hours. ATAC-seq analysis was performed as two biological replicates, which were highly similar, and a single replicate for each condition is shown. The number of accessible peaks in control i-SOX2-ONS76 cells averaged approximately 177,500 peaks.

### Differential effects of SOX2 and SOX2(G76P) at the gene level

Besides substantial differences between the effects of SOX2 and SOX2(G76P) at the genomic level, there are significant differences between the effects of SOX2 and SOX2(G76P) on the expression of individual genes. RNA-Seq data indicated that elevating SOX2 upregulated genes, such as *NRCAM, NRP2* and *SOX2-OT*, that were not upregulated by SOX2(G76P) (Fig. 4A; Supplemental Fig. S5A, S6A). Conversely, there were genes, such as *CPA5, SCIRT, and POU2F2* that were upregulated by SOX2(G76P), but not SOX2 (Fig. 5A). Examination of the SOX2 ChIP-seq profiles of these genes reflects the differential action of SOX2 and SOX2(G76P). In the case of *NRCAM*, which was only upregulated by SOX2 (>30 fold), there were substantial increases in SOX2 binding within the NRCAM gene locus, whereas there were no significant changes in its ChIP-seq gene profile when SOX2(G76P) was elevated (Fig. 4B). Not surprisingly, regions bound by SOX2 within the *NRCAM* gene were accompanied by increases in ATAC-seq peaks, that were not observed in the SOX2(G76P) elevated cells (Fig. 4B). A similar pattern was observed for *NRP2*, which was upregulated ∼2-fold by SOX2, but not by SOX2(G76P) (Supplemental Fig. S5A). The increased expression of *NRP2* corresponded to enhanced SOX2 binding, which was confirmed by ChIP-qPCR analysis of two gene loci (Supplemental Fig. S5B), as well as by SOX2 ChIP-seq analysis and increased chromatin accessibility (Supplemental Fig. S5C). These results were not observed with SOX2(G76P) (Supplemental Fig. S5B,C). Interestingly, within the gene locus of *SOX2-OT*, which was strongly upregulated by SOX2 (>80-fold) (Supplemental Fig. S6A), there were numerous new small SOX2 peaks in the SOX2-elevated cells that were not observed in SOX2(G76P) elevated cells (Supplemental Fig. S6B). We confirmed the differential association of SOX2 and SOX2(G76P) within the *SOX2-OT* gene by performing ChIP-qPCR utilizing primer sets that align within two of the ChIP-seq peaks (Supplemental Fig. S6C). At both of these regions, we observed large increases (>30-fold) in SOX2 association, but little or no enrichment or change in chromatin accessibility when SOX2(G76P) was expressed.

**Figure 4.**
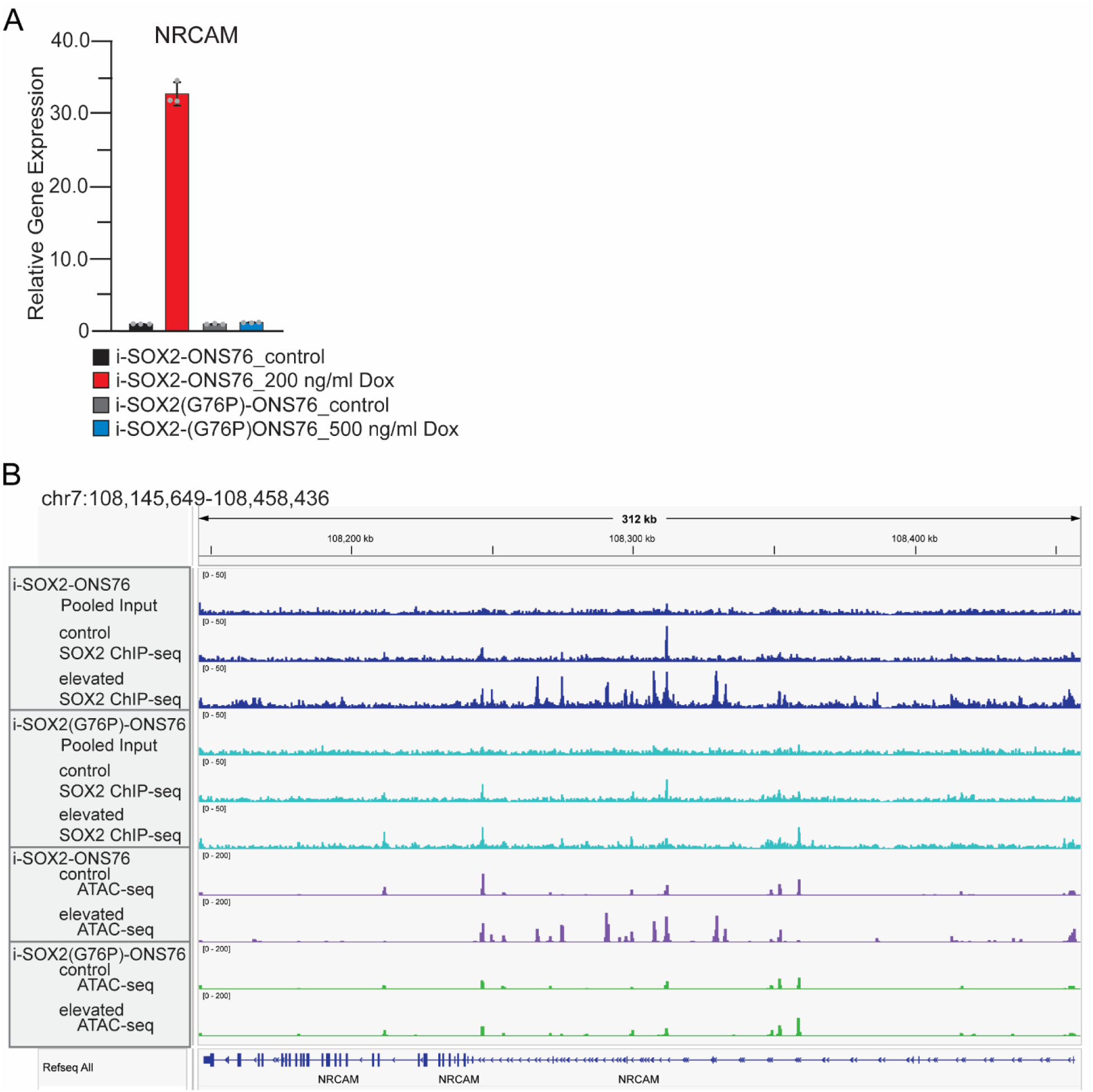
Changes in expression, SOX2 binding, and chromatin accessibility of *NRCAM*. (A) RT-qPCR analysis of *NRCAM* expression relative to *GAPDH* using samples processed from i-SOX2-ONS76 and i-SOX2(G76P)-ONS76 cells cultured in the absence or presence of Dox for 48 hours. Gray dots represent replicates, and error bars represent standard deviation. (B) IGV screen shots of bigWig peaks at the *NRCAM* gene identified through SOX2 ChIP-seq and ATAC-seq experiments from i-SOX2-ONS76 and i-SOX2(G76P)-ONS76 control cells and cells cultured for 48 hours with Dox.

**Figure 5.**
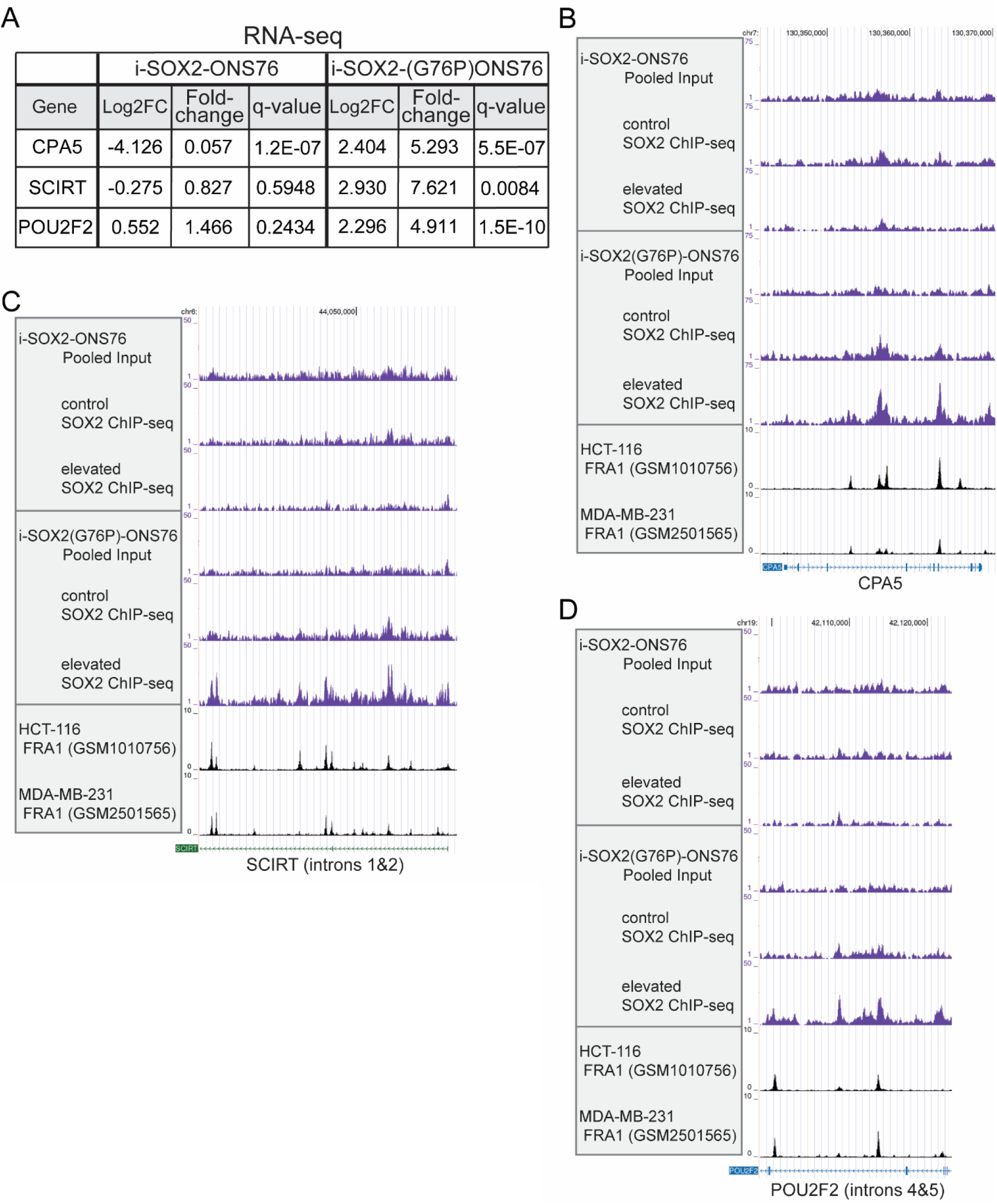
SOX2 and SOX2(G76P) ChIP-seq analysis aligned with FRA1 ChIP-seq for specific genes. (A) Differential expression of control and Dox-induced i-SOX2-ONS76 and i-SOX2(G76P)-ONS76 cell RNA-seq data for the genes: *CPA5*, *SCIRT*, and *POU2F2*. Log2FC is Log2 fold change. (B-D) UCSC Genome Browser screen shots of bigWig peaks identified through our SOX2 ChIP-seq data collected from i-SOX2-ONS76 and i-SOX2(G76P)-ONS76 control cells and cells cultured with Dox for 48 hours (elevated) that overlapped with FRA1 binding in HCT-116 and MDA-MB-231 from published datasets within (B) the *CPA5* gene, (C) introns 1 and 2 of the *SCIRT* long non-coding RNA, and (D) introns 4 and 5 of the *POU2F2* gene.

We also observed prominent differences in the ChIP-seq profiles of SOX2 and SOX2(G76P) at genes only upregulated by SOX2(G76P) (Fig. 5). In the cases of *CPA5, SCIRT, and POU2F2*, which were each upregulated approximately 5-7-fold by SOX2(G76P), but not by SOX2 (Fig. 5A), there were substantial increases in ChIP-seq peaks when SOX2(G76P) was elevated, but not when SOX2 was elevated (Fig. 5B-D). Peaks within these three genes clustered into ChIP-seq cluster 1 (*CPA5* and *SCIRT*) and cluster 4 (*SCIRT* and *POU2F2*) (Fig. 2B). Interestingly, cluster 1 lacks SOX motifs entirely and cluster 4’s top 15 motifs primarily consist of AP-1 family motifs. This raised the possibility that SOX2(G76P) could associate with these genes indirectly by associating with other transcriptional machinery without itself binding directly to DNA. One likely candidate for the indirect association of SOX2(G76P) with these genes is the transcription factor FRA1, which was the top ranked motif for both clusters 1 and 4 (Fig. 2C). Significantly, comparison of the SOX2(G76P) ChIP-seq data profiles of *CPA5, SCIRT, and POU2F2* with previously published FRA1 ChIP-seq data generated in HCT116 cells and MDA-MB-231 cells (Myers 2012; Gallenne et al. 2017), revealed regions of FRA1 peaks overlapping with SOX2(G76P) peaks for each of these genes (Fig. 5B-D).

The significance of the overlap between SOX2(G76P) and FRA1 peaks at *CPA5, SCIRT, and POU2F2* is highlighted by the overlap between SOX2(G76P) and FRA1 peaks at the genome level. Importantly, FRA1 ChIP-seq peaks found in HCT116 cells and MDA-MB-231 cells (Myers 2012; Gallenne et al. 2017) overlap with SOX2(G76P) ChIP-seq peaks present at many genes in cluster 1, 2 and 4 from the Dox-induced i-SOX2(G76P)-ONS76 cells (Fig. 6). Significantly, in contrast to the overlap between SOX2(G76P) and FRA1, no overlap was observed between SOX2(G76P) and EZH2 (Fig. 6). Moreover, there is no overlap in clusters 3 and 5, which primarily consist of targets containing SOX motifs (Fig. 2C). Collectively, these findings and those discussed above indicate that SOX2 and SOX2(G76P) not only associate with different gene loci they also regulate those genes they associate with. Equally significant, our findings support the conclusion that SOX2(G76P) associates with chromatin indirectly by associating with other chromatin bound proteins.

**Figure 6.**
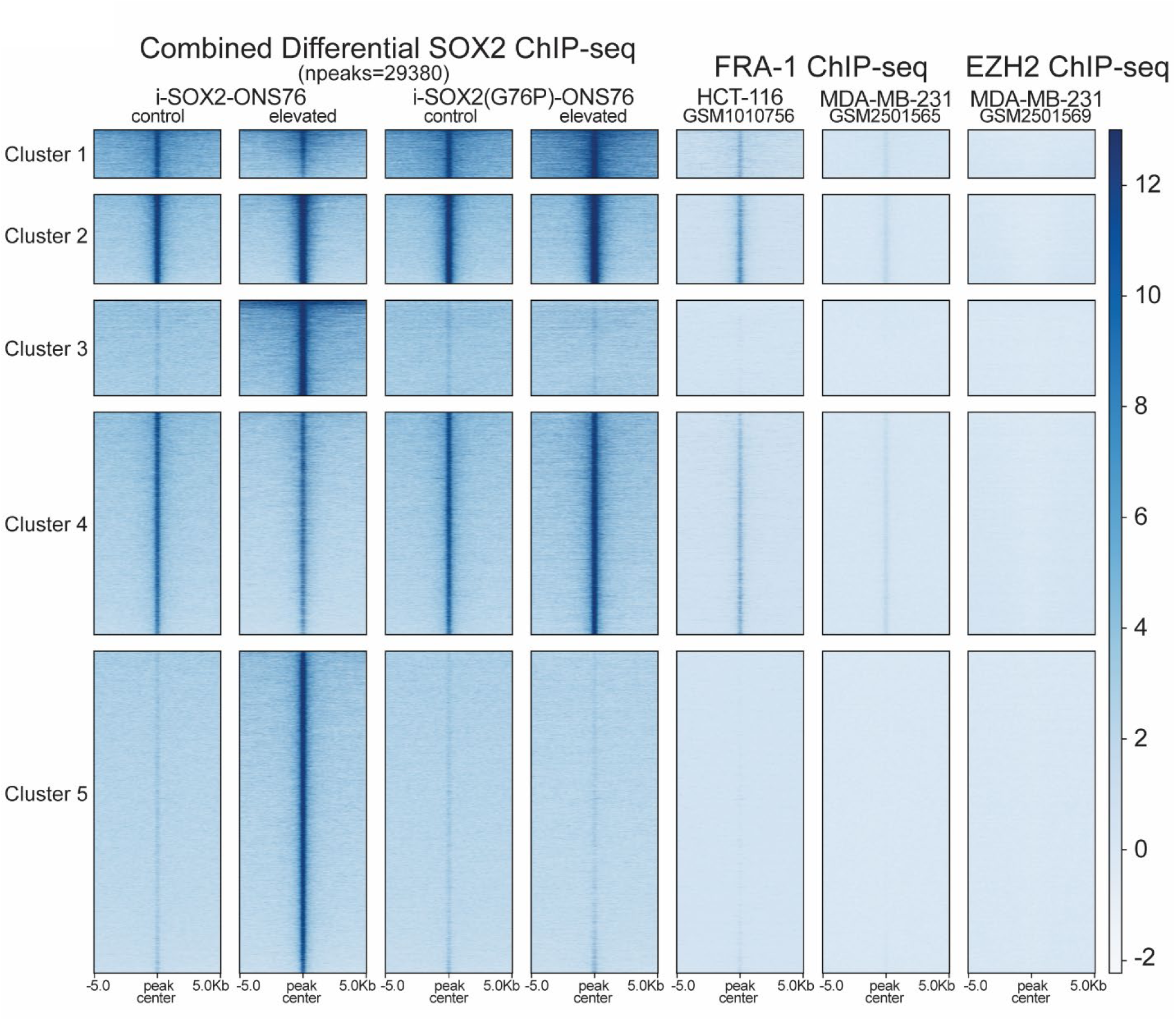
Unsupervised clustering of combined differential SOX2 ChIP-seq genomic peaks integrated with published FRA1 ChIP-seq datasets. Heatmap of clustered genomic regions using a combined differential of SOX2 and SOX2(G76P) ChIP-seq profiles with control i-SOX2-ONS76 cells as the reference for the comparison of SOX2 ChIP-seq from 48-hour control and Dox-induced i-SOX2-ONS76 cells and control and Dox-induced i-SOX2(G76P)-ONS76 cells, which were integrated with FRA1 ChIP-seq from two different cell lines: HCT-116 and MDA-MB-231 cells (GSM1010756 and GSM2501565, respectively). EZH2 ChIP-seq from MDA-MB-231 cells (GSM2501569) served as a negative control.

### SOX2 and SOX2(G76P) associate with a small subset of the same gene loci

Although it is evident that SOX2 and SOX2(G76P) largely associate with different DNA sequences, multiple lines of evidence indicate that SOX2 and SOX2(G76P) can associate with a small subset of the same gene sequences genome-wide, including those lacking SOX2 motifs. Examination of DNA motifs in clusters 1 and 4 (Fig. 2B,C) indicates that there are a large set of peaks enriched in FRA1 (AP-1) motifs that are shared by SOX2 and SOX2(G76P).

Similarly, peaks shared by SOX2 and SOX2(G76P) in cluster 2 are enriched in multiple ETS motifs (Fig. 2B,C). Overlap between SOX2 and SOX2(G76P) is also evident at the gene level. Screenshots of ChIP-seq peaks present in three genes, *GADD45A, WDR74, and PRKAG2*, show significant overlap between SOX2 and SOX2(G76P) in both control and Dox-induced i-SOX2-ONS76 and i-SOX2(G76P)-ONS76 cells, in addition to overlapping with FRA1 peaks previously observed in HCT-116 and MDA-MB-231 cells (Supplemental Fig. S7).

Deeper analysis of our ChIP-seq data provides further support for association of both SOX2 and SOX2(G76P) at some of the same genomic regions lacking SOX motifs. The ChIP-seq analysis in Figure 2 compared the association of SOX2 and SOX2(G76P) with gene loci using a composite list of peaks associated with SOX2 and/or SOX2(G76P). Thus, that analysis did not differentiate between peaks that are associated only with SOX2, only with SOX2(G76P), or both SOX2 and SOX2(G76P). However, by reanalyzing the ChIP-seq data and focusing our analysis only on the subset of SOX2(G76P) ChIP-seq peaks that change when SOX2(G76P) was expressed in the Dox induced i-SOX2(G76P)-ONS76 cells, we observed many of the same ChIP-seq peaks for SOX2 in the control and the Dox induced i-SOX2-ONS76 cells (Fig. 7).

**Figure 7.**
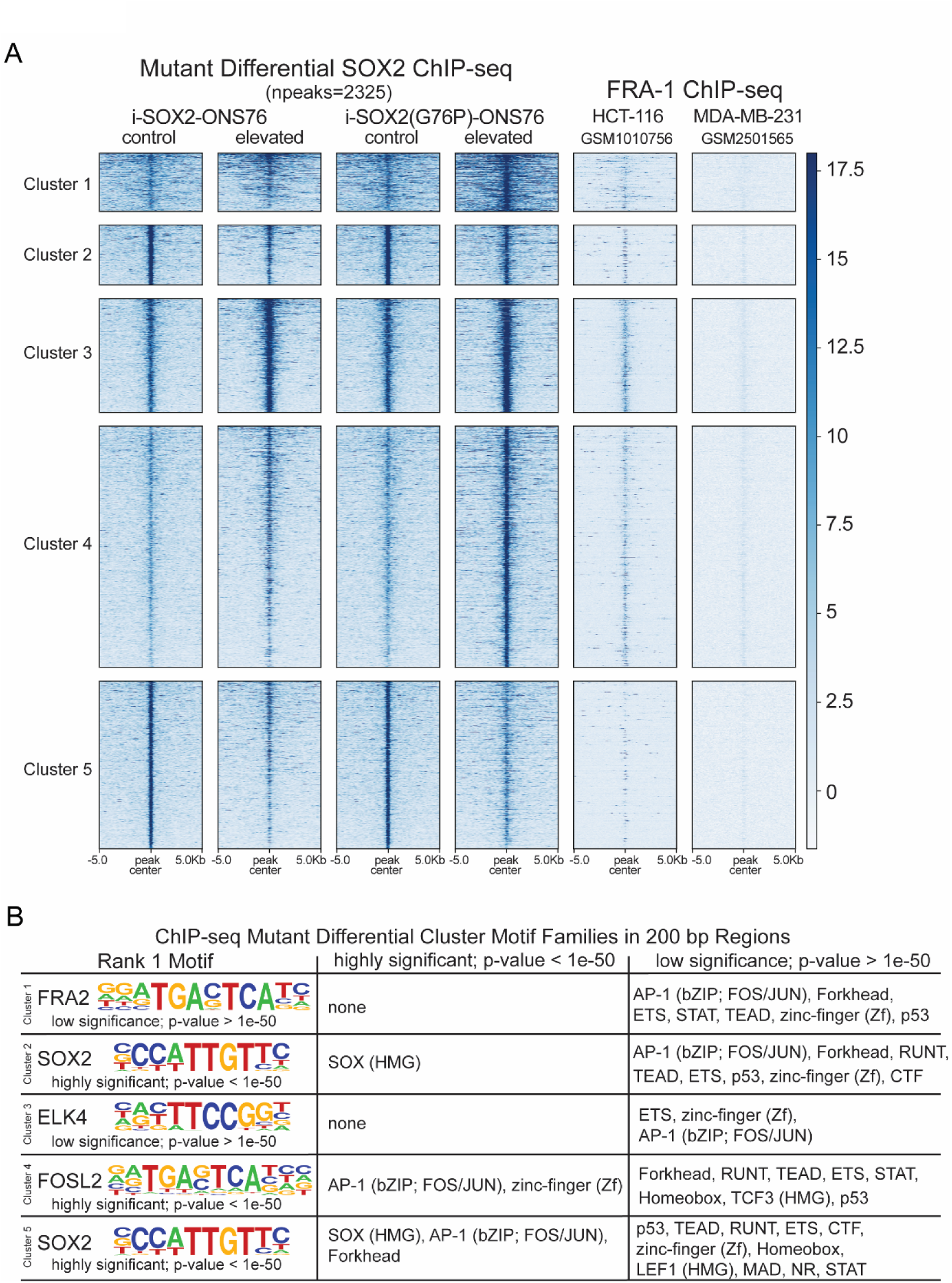
Unsupervised clustering of SOX2(G76P) differential SOX2 ChIP-seq genomic peaks integrated with published FRA1 ChIP-seq dataset. (A) Heatmap of clustered genomic regions using only peaks that exhibit differential SOX2(G76P) ChIP-seq profiles with control i-SOX2-ONS76 cells as the reference to scale the comparison of SOX2 ChIP-seq from 48-hour control and Dox-induced i-SOX2-ONS76 cells and control and Dox-induced i-SOX2(G76P)-ONS76 cells, which were integrated with FRA1 ChIP-seq from two different cell lines: HCT-116 and MDA-MB-231 cells (GSM1010756 and GSM2501565, respectively). (B) The top motif as well as other highly significant (p-value < 1e-50) and low significance (p-value > 1e-50) motif families identified using HOMER in 200 bp regions within each of the ChIP-seq clusters.

Moreover, the increases in peaks in clusters 1, 3, and 4 when SOX2(G76P) was expressed matched the increases in peaks when SOX2 was elevated (Fig. 7). Importantly, peaks in cluster 4 were enriched for FOSL2 (AP-1) and several other statistically significant motifs (p<1e-50), but completely devoid of SOX motifs (Supplemental Table S1). Additionally, we observed overlapping peaks from previously published FRA1 ChIP-seq profiles (Myers 2012; Gallenne et al. 2017) in clusters 3 and 4 (Fig. 7A). Taken together, these findings support two conclusions: SOX2 and SOX2(G76P) associate with a small set of gene sequences in common, and SOX2, like SOX2(G76P), can associate with this small set of genes indirectly through its association with other chromatin-associated transcriptional machinery, rather than binding directly to DNA at SOX motifs.

### SOX2 and SOX2(G76P) behave differently in CUT&RUN

Given that SOX2 associates with most genes by binding to SOX2 motifs, whereas SOX2(G76P) does not, we tested the possibility that their chromatin association at most genes might be topologically different from one another. To test this premise, we chose to perform CUT&RUN because it requires that micrococcal nuclease (MNase) fused to protein A and/or G, which is bound to the SOX2 antibody, is positioned close enough to cleave DNA on both sides of the chromatin-associated protein complex (Skene and Henikoff 2017). In contrast, ChIP-seq does not depend on proximity of the chromatin-bound protein to DNA because sonication is used to fragment formaldehyde cross-linked DNA:protein complexes (DeCaprio and Kohl 2020; Ormsbee Golden et al. 2024). Using control and Dox induced i-SOX2-ONS76 and control and Dox induced i-SOX2(G76P)-ONS76 cells, we observed remarkably different results when SOX2 and SOX2(G76P) CUT&RUN were performed. Unsupervised clustering demonstrated that elevation of SOX2 dramatically increased its association with genomic regions within five clusters, especially within clusters 1, 2 and 4 (Fig. 8A). Not surprisingly, SOX sequences were the most common motifs bound by SOX2 in these three genomic clusters (Fig. 8B). In strong contrast, there were relatively few peaks in genomic clusters 1-5 when SOX2(G76P) was induced in i-SOX2(G76P)-ONS76 cells. This lack of pronounced SOX2(G76P) enrichment is also evident when comparing the profiles of entire chromosomes, such as chromosome 7 (Supplemental Fig. S8). To validate these findings, we repeated the CUT&RUN analysis using a different SOX2 antibody. Specifically, we used the same SOX2 antibody used for the ChIP-seq studies (Fig. 2). Importantly, the same results were obtained (Supplemental Fig. S9). With both SOX2 antibodies, there was a substantial increase in the number of CUT&RUN peaks when SOX2 was elevated; whereas there was little if any increase in the number of CUT&RUN peaks when SOX2(G76P) was expressed. These findings provide substantial evidence that SOX2 and SOX2(G76P) associate with chromatin in significantly different ways. If SOX2(G76P) were bound directly to DNA, we should have observed more prominent increases in the CUT&RUN profile when SOX2(G76P) was expressed.

**Figure 8.**
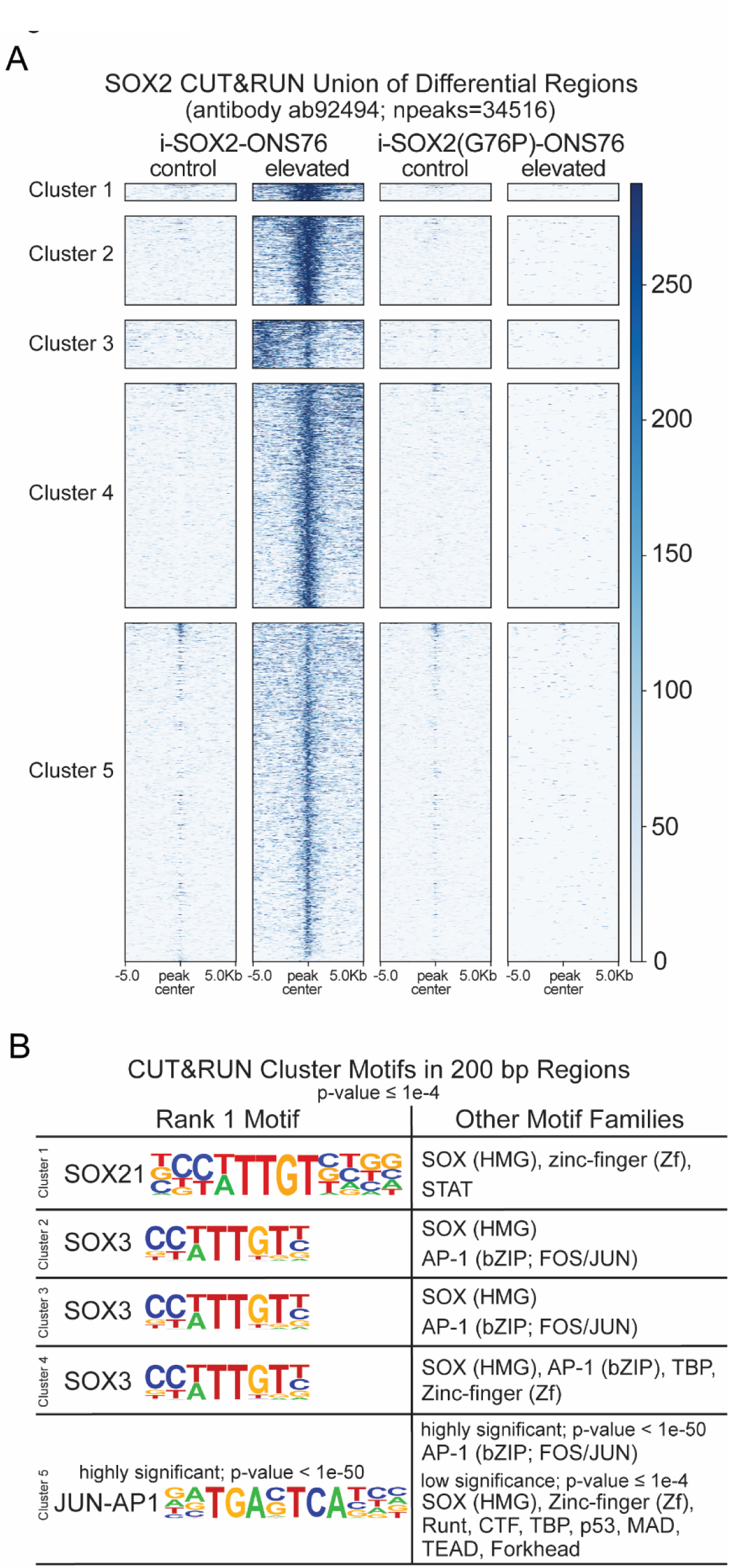
SOX2 and SOX2(G76P) CUT&RUN analysis. (A) Heatmap of clustered genomic sequences from the combined differential of SOX2 and SOX2(G76P) CUT&RUN results with the control i-SOX2-ONS76 cell results as the reference for comparison of SOX2 CUT&RUN from 48-hour control and Dox-induced (elevated) i-SOX2-ONS76 cells, and control or Dox-induced (elevated) i-SOX2(G76P)-ONS76 cells. CUT&RUN was performed as two replicates, which were similar. Only one replicate for each condition is shown. (B) Top ranked known motif as well as other motif families within each CUT&RUN cluster identified using HOMER in 200 bp regions.

## Discussion

In this study, we created a SOX2 DNA binding mutant, SOX2(G76P), and compared its function head-to-head with SOX2. We demonstrate that the *in vitro* binding of this mutant to DNA is impaired, yet it is biologically functional. Like SOX2 (Metz et al. 2020b), SOX2(G76P) inhibits proliferation of ONS76 cells in a dose-dependent manner, and like SOX2 (Metz et al. 2022) it can dramatically reprogram the transcriptome when expressed in their respective Dox-induced cells. However, the effects of SOX2(G76P) on the transcriptome differ substantially from those of SOX2. Gene Ontology analysis indicates that SOX2 and SOX2(G76P) upregulate and downregulate very different biological processes.

Fully consistent with the differential effects on the transcriptome, ChIP-seq data indicates that SOX2 and SOX2(G76P) associate most prevalently with different genomic sequences in the Dox-induced cells. The importance of the contrasting results is highlighted by differences in the DNA motifs present in the genomic sequences associated with SOX2 and SOX2(G76P). SOX motifs predominated in the genomic clusters 3 and 5 that exhibited increased SOX2 enrichment when SOX2 was elevated in i-SOX2-ONS76 cells (Fig. 2). This is consistent with numerous earlier reports that SOX2 binds to SOX motifs (Harley et al. 1994; Kamachi et al. 1999; Boyer et al. 2005). In contrast, Fra1 is the top ranked motif in genomic clusters 1 and 4, which only exhibited increased enrichment when SOX2(G76P) was expressed in Dox-induced i-SOX2(G76P)-ONS76 cells (Fig 2). Furthermore, none of the high confidence motifs in cluster 1 are SOX motifs. Important differences between SOX2 and SOX2(G76P) are also evident when their effects on chromatin accessibility are compared. ATAC-seq data indicates that elevating SOX2 leads to large increases in the number of chromatin accessible peaks: ∼30% above basal levels. In contrast, there were very modest increases in the number of chromatin-accessible peaks, ∼2.5%, when SOX2(G76P) was expressed in the Dox-induced cells. Collectively, these findings indicate that both SOX2 and SOX2(G76P) can reprogram the transcriptome and associate with chromatin, but they do so in significantly different ways.

We also identified significant differences between SOX2 and SOX2(G76P) at the single gene level. Consistent with our ChIP-seq, RNA-Seq, and ATAC-seq data, we show *NRCAM*, *NRP2*, and *SOX2-OT* are each upregulated by SOX2, but not by SOX2(G76P), whereas *CPA5, SCIRT, and POU2F2* are upregulated when SOX2(G76P), but not SOX2, is expressed in their respective Dox-induced cells. The upregulation of *SOX2-OT* by SOX2 is noteworthy because the *SOX2* gene is located within intron seven of *SOX2-OT* and because previous studies have shown that *SOX2-OT* can upregulate *SOX2* expression (Askarian-Amiri et al. 2014). Thus, a positive-feedback loop appears to exist between *SOX2* and *SOX2-OT* that is likely to depend on direct binding of SOX2 to the *SOX2-OT* gene. Together, the differential effects on the expression of multiple genes illustrate that significant differences between SOX2 and SOX2(G76P) are not only evident at the genome level, but also at the single gene level.

The apparent lack of SOX motifs in the majority of genomic sequences associated with SOX2(G76P) leads us to conclude that, unlike SOX2, which primarily associates with chromatin by binding to DNA, SOX2(G76P) associates primarily, if not exclusively, with chromatin indirectly as part of protein complexes containing other transcription factors. We refer to this as a “piggyback” mechanism for regulating gene transcription. While it is conceivable that SOX2(G76P) can bind directly to DNA at non-SOX motifs, for example, at FRA1 (AP-1), ELK, and ATF3 motifs, it is far more likely that it associates indirectly with genes as part of larger protein complexes. Importantly, we demonstrated previously that SOX2 in nuclear extracts is present in large protein complexes, some larger than 880 kDa (Cox et al. 2010). Moreover, proteomic analysis of the SOX2 interactome indicates that SOX2 associates with a wide array of transcription factors and transcriptional machinery in embryonic stem cells, medulloblastoma, and glioblastoma cells (Mallanna et al. 2010; Gao et al. 2012; Cox et al. 2013). This includes protein complexes containing both SOX2 and the AP-1 protein JUND in medulloblastoma cells (Cox et al. 2013). Equally important, there are clear precedents of transcription factors associating both directly and indirectly with chromatin. It is widely recognized that glucocorticoid receptors bind directly to glucocorticoid response elements and indirectly to other gene regulatory sequences as part of protein complexes containing other DNA binding transcription factors (Miranda et al. 2013). This is also true for estrogen receptors. Early work established that estrogen receptors, which typically bind to estrogen receptor elements, also associate indirectly with chromatin by protein-protein tethering to AP-1 (Jakacka et al. 2001; Carroll et al. 2006).

The conclusion that SOX2(G76P) associates with chromatin indirectly is further supported by the dramatic differences in the CUT&RUN results observed for SOX2 and SOX2(G76P). Elevation of SOX2 in the Dox-induced i-SOX2-ONS76 cells led to substantial increases in enrichment within all five CUT&RUN clusters (Fig. 8A). In strong contrast, expression of SOX2(G76P) in their Dox-induced cells led to no substantial increases in any of the five CUT&RUN clusters. These results are in stark contrast to the large quantities of both SOX2 and SOX2(G76P) protein that were extracted from chromatin isolated from their respective Dox-induced cells (Fig S3A). Moreover, we observed large increases in the number of ChIP-seq peaks in both the Dox-induced i-SOX2-ONS76 cells and the Dox induced i-SOX2(G76P)-ONS76 cells (Fig 2A). We suggest that the contrasting outcomes between the CUT&RUN and ChIP-seq results are due to fundamental differences in their methodologies, as discussed earlier, and illustrate significant differences in the proximity of SOX2 and SOX2(G76P) to DNA when associated with chromatin. Taken together, our CUT&RUN and ChIP-seq results provide strong support for the conclusion that the vast majority, if not all, of SOX2(G76P)-chromatin associations occur indirectly by a piggyback mechanism. Importantly, our findings with SOX2(G76P) also lend further support for the emerging view that the gene target specificity of transcription factors *in vivo* is not only influenced by their DBD, but also by other regions of transcription factors, especially their IDRs (Brodsky et al. 2020; Jonas et al. 2025; Abidi et al. 2026). Equally important, the differential results obtained with ChIP-seq and CUT&RUN suggest that CUT&RUN may fail to adequately detect transcription factors that associate with chromatin indirectly.

Finally, we have provided several lines of evidence that SOX2 and SOX2(G76P) can associate with chromatin at some of the same gene loci without directly binding to DNA. Importantly, our ChIP-seq data indicates that there are genomic sequences bound by SOX2 that lack apparent SOX motifs. In both control and Dox induced i-SOX2-ONS76 cells, SOX2 associates with the gene loci in cluster 1 (Fig. 2B), where the top ranked motifs of this cluster are FRA1 and other AP-1 motifs, without high confidence SOX motifs (Fig. 2C). Secondly, by focusing on the small set of SOX2(G76P) differentially enriched peaks that were identified in ChIP-seq (Fig. 7), we show that SOX2 association increases within the gene sequences in clusters 1, 3, and 4 in both Dox induced i-SOX2-ONS76 and Dox-induced i-SOX2(G76P)-ONS76 cells, where there is also overlap with FRA1 ChIP-seq peaks. Importantly, cluster 4 contains primarily AP-1 family motifs within the high confidence motifs identified, as well as a range of low confidence motifs, including Forkhead, ETS, and other bZIP family members, but no SOX motifs. Moreover, there are no low or high-confidence SOX motifs in clusters 1 or 3 (Fig. 7). Additionally, the gene profiles for *GADD45A, WDR74,* and *PRKAG2* show overlapping ChIP-seq peaks for both SOX2(G76P) and SOX2, that coincide with previously identified ChIP-seq FRA1 peaks identified in HCT-116 cells and MDA-MB-231 cells (Supplemental Fig. S7). Importantly, if SOX2 is not associating with these gene loci indirectly, we would need to conclude that SOX2 can bind to DNA at non-SOX motifs. While this is conceivable, we believe indirect chromatin association at some gene loci is far more likely. Collectively, our findings provide a new perspective on SOX2 and lead to an important conclusion. While SOX2 primarily associates with genomic sequences directly by binding to DNA (SOX motifs), it can also associate indirectly with a subset of gene loci through its association with other chromatin bound transcriptional machinery. It is likely that the indirect association of SOX2 with a subset of genomic sequences has not been previously recognized due to the prevalence of SOX2 directly binding at SOX motifs genome-wide. The importance of indirect chromatin association is more apparent in our study with SOX2(G76P) because it is unable to bind at SOX motifs but maintains its ability to associate with other transcription factors for indirect binding at their DNA binding motifs.

In conclusion, we provide multiple lines of evidence that SOX2(G76P) associates with chromatin indirectly by a “piggyback” mechanism through its association with other chromatin-associated proteins, including AP-1 complexes. Importantly, this study also provides new and important insights into molecular mechanisms through which the stem cell transcription factor SOX2 regulates the transcription of its target genes. Specifically, the findings presented lead to two important conclusions. First, selection of gene targets by SOX2 is not solely determined by its DNA binding domain. Second, SOX2 not only associates with chromatin directly by binding to SOX motifs but can also associate with a subset of gene loci indirectly through its association with other chromatin-associated proteins.

## Materials and Methods

### Site-directed Mutagenesis and Plasmid Production

The mouse Flag-Sox2-CMV5 expression plasmid created previously (Wiebe et al. 2003) and the puromycin-selectable human Flag-Strep-tagged SOX2 lentiviral vector, pLVX-tetO-(fs)SOX2 (Cox et al. 2012), were altered through site-directed mutagenesis to introduce a point mutation (mouse G78P, human G76P) to disrupt the DBD of SOX2. Site-directed mutagensis was accomplished using a modification of the QuikChange method (Stratagene, La Jolla, CA). The primer used to modify human SOX2 with its reverse complement was 5’-GAGATCAGCAAGCGCCTGCCTGCAGAGTGGAAACTTTTGTC-3’. The site-directed mutagenesis also created a diagnostic PstI site (CTGCAG) in the Sox2 sequence at human codon 77 (mouse codon 79), which was a silent mutation of the alanine codon GCA from human GCC and mouse GCG. There was no other difference between the human and mouse sequences in the stretch covered by these primers. The two DNA-binding mutant SOX2 plasmids were sequenced by the UNMC DNA Sequencing Core Facility to confirm that the desired changes in the SOX2 sequence had been introduced. These methods led to the production of plasmids referred to in this study as Flag-Sox2(G78P)-CMV5 expression plasmid and pLVX-tetO-(fs)SOX2(G76P) lentiviral plasmid.

### Lentivirus Production

Two separate viral stocks were produced, one of pLVX-tetO-(fs)SOX2(G76P) (plasmid described above) and the other of the reverse tetracycline transactivator M2rtTA (Addgene, Plasmid 20342). The lentiviral production, particle filtration, and collection steps have been described previously (Cox et al. 2011).

### Electrophoretic Mobility Shift Assay (EMSA)

Transient transfection of 1.5 x 10^6^ 293T cells in 100 mm dishes with expression plasmids was accomplished using the calcium phosphate precipitation method (Ma et al. 1992), followed by nuclear protein isolation (as described below). Expression plasmids for mouse Flag-Sox2-CMV5 and mouse Flag-Sox2(G78P)-CMV5 (described above) were used to conduct EMSA as detailed previously [22]. The forward and reverse HDAC2 probe were 5’-TAGCCACTTCTTTGTTGCTCTTTGTTCGATC-3’ and 5’-TAGCGATCGAACAAAGAGCAACAAAGAAGTG-3’, respectively (Fig S1). The antibody used for this experiment was anti-Flag (Sigma, F1804).

### Cell Culture

ONS76 medulloblastoma cells capable of inducible expression of a DNA-binding mutant for SOX2 were engineered utilizing the lentiviral particles described above in the transduction and selection process described previously (Wuebben et al. 2016; Metz and Rizzino 2019). These cells, referred to as i-SOX2(G76P)-ONS76, and their previously described wild-type counterpart, i-SOX2-ONS76 cells (Metz et al. 2020b), were cultured in Dulbecco’s Modified Eagle Medium (12100046, Life Technologies) supplemented with 10% HyClone Characterized Fetal Bovine Serum (SH30910.03, Cytiva, Fisher Scientific) and 1% Pen-Strep (P3032-10MU & S6137-25G, SIGMA). Both cell lines were maintained at 37°C and 5% CO2. To induce expression of either the wild-type SOX2 or DNA-binding mutant SOX2(G76P) in the i-SOX2-ONS76 or i-SOX2(G76P)-ONS76 cells, respectively, Doxycycline (Dox) (TA9H30375FFD, SIGMA), suspended in water, was premixed in the media used to replenish cells after an initial 24 hours of culture (control cells were replenished with plain complete media), and then cultured for another 48 hours. Unless otherwise noted, the pre-mixed Dox media for i-SOX2-ONS76 cells was brought to a final concentration of 200 ng/ml Dox and i-SOX2(G76P)-ONS76 to 500 ng/ml, which were the concentrations found to result in 40-50% growth inhibition of each cell line (Metz et al. 2020b) (Fig 1B). The proliferation of i-SOX2-ONS76 cells following Dox induction of elevated SOX2 was previously assessed (Ormsbee Golden et al. 2024), and following those methods, we similarly determined the proliferation of i-SOX2(G76P)-ONS76 cells by performing MTT assays with triplicate samples (Fig 1B). Cells tested negative for mycoplasma using a Myco Sensor PCR Assay Kit (302109, Agilent). Cell line authenticity of i-SOX2-ONS76 was performed previously (Metz et al. 2022) and was confirmed in the same manner for i-SOX2(G76P)-ONS76.

### Protein Isolation and Western Blotting

Whole cell protein extracts were prepared from i-SOX2(G76P)-ONS76 cells. After 24 hours of culture, one control plate was replenished with fresh complete media, and the other plates were replenished with media pre-mixed with Dox at the concentrations noted. Isolation of whole cell protein was conducted following the steps described in the Santa Cruz western blotting RIPA buffer monolayer cell processing protocol using 300 μl RIPA buffer (150 mM NaCl, 50 mM Tris-HCl pH 7.4, 1% IGEPAL, 0.25% Sodium Deoxycholate, and 1 mM EDTA) supplemented with the protease inhibitors detailed in the Supplemental Materials.

Chromatin-bound proteins were isolated from cells cultured the same as above and after 48 hours, protein extracts from control and Dox-induced i-SOX2-ONS76 and i-SOX2(G76P)-ONS76 cells were prepared using the Pierce NE-PER™ nuclear and cytoplasmic extraction kit (78833, Pierce, Thermo Fisher Scientific), as described previously (Kopp et al. 2008; Ormsbee Golden et al. 2024). However, the NE-PER™ kit does not utilize the insoluble fraction that remains after the nuclear extract is isolated from lysed nuclei, and the residual insoluble fraction consists of chromatin. The chromatin-bound proteins within this fraction were isolated using Benzonase following a previously established protocol (Gillotin 2018) that we modified to avoid excessive dilution of the protein. The details of our modification to this protocol and the remaining steps for all protein concentration measurements and western blot reagents are described in the Supplemental Materials.

The primary antibodies used were SOX2 (Cell Signaling Technology [CST], 3579S, 1:1000); HDAC1 (CST, 34589S, 1:1000); and β-TUBULIN (CST, 2146S, 1:1000); and the secondary anti-rabbit antibody was from CST (7054S, 1:5000).

### RNA isolation, cDNA synthesis, RT-qPCR and RNA-seq

RNA isolation, cDNA synthesis, and RT-qPCR were performed as described previously (Ormsbee Golden et al. 2024). Primers used for RT-qPCR are provided in Supplemental Table S2. Primers against GAPDH were utilized for normalization in the RT-qPCR analysis and relative gene expression was characterized by the comparative CT method (2^-ΔΔCT). For RNA-Seq, RNA was isolated from i-SOX2(G76P)-ONS76 cells as described previously (Metz et al. 2022). The cells were cultured for a total of three days and for the last 48 hours, the cells were cultured in the presence or absence of Dox. Library preparation and transcriptome sequencing on an Illumina NovaSeq 6000 platform were performed by Novogene (Beijing, China) and additional RNA-seq analysis details are provided in the Supplemental Materials.

### ChIP and ChIP-qPCR

ChIP-qPCR was performed as described previously (Ormsbee Golden et al. 2024). For these studies, the following antibodies were used: RNA Pol II (Millipore, 05-623, 1:50), p300 (CST, 54062, 1:50), BRD4 (CST, 13440, 1:50), H3K27ac (CST, 8173, 1:100), H3K27me3 (CST 9733S, 1:50), SOX2 (Active Motif 39823, 1:50). The primers used for ChIP-qPCR are listed in Supplemental Table S2.

### Chromatin Immunoprecipitation Sequencing (ChIP-seq)

ChIP-seq was performed in duplicate (as biological replicates) for each condition and cell line through services obtained from Active Motif using their provided SOX2 antibody (Active Motif, cat# 39823). In preparation, cells seeded at 1.0 x 10^6^ each in four 100 mm plates for both cell lines, i-SOX2-ONS76 and i-SOX2(G76P)-ONS76 (eight plates total), were cultured for an initial 24 hours and then replenished with control complete media or pre-mixed Dox media. After 48 hours of Dox exposure, the cells were prepared as directed by Active Motif’s ChIP Cell Standard Sample Preparation Protocol, which is described in detail, along with library preparation and initial data analysis, in the Supplemental Materials.

### Assay for Transposase-Accessible Chromatin using sequencing (ATAC-seq)

Cells were seeded in 60 mm plates at 0.2 x 10^6^ for each of i-SOX2-ONS76 in four plates and i-SOX2(G76P)-ONS76 in four plates to generate control and Dox-induced duplicates (biological replicates). Cells were released from attachment with trypsin and counted to prepare 100,000 fresh cell aliquots in 1.5 ml centrifuge tubes to follow the ATAC-seq kit protocol from Active Motif (53150). The kit includes all the necessary buffers to isolate nuclei and perform the tagmentation reaction with the provided assembled transposomes, as well as prepare libraries. Details for library preparation, quantification, sequencing, and peak calling are provided in the Supplemental Materials.

### Cleavage Under Targets & Release Using Nuclease (CUT&RUN)

Preparation of cells for CUT&RUN involved seeding of each cell line, i-SOX2-ONS76 and i-SOX2(G76P)-ONS76, in four 100 mm plates at 1.3×10^6^ each. After 24 hours of culture, the media for each plate was replenished to provide two control plates of each cell line and two pre-mixed Dox media plates. After 48 hours of Dox-induced SOX2 expression, CUT&RUN was performed as described previously (Brahma and Henikoff 2019, 2022), first with the CUT&RUN validated SOX2 antibody from Abcam (ab92494) (Fig 8) in duplicate, followed by a repeat of CUT&RUN in duplicate using the ChIP-seq antibody (Active Motif, cat# 39823) (Supplemental Fig S9) that we used for our SOX2 ChIP-seq studies. Further details for our CUT&RUN sample processing, library preparation, and peak calling are provided in the Supplemental Materials.

### Library Data Analysis and Visualization

For visualization of peaks from CUT&RUN, ATAC-seq, and ChIP-seq experiments, bigWig files were generated from .bam files using deepTools (Ramírez et al. 2016) bamCoverage and peak heatmaps were generated using deepTools computematrix (referencepoint=center and binsize=50) and plotHeatmap functions. The subset of differential peaks for most analyses was identified for each group [i-SOX2-ONS76 Dox-induced vs control and i-SOX2(G76P)-ONS76 Dox-induced vs control] and merged to create a list of genomic regions for visualization. The subset of differential peaks for Figure 7 identified for i-SOX2(G76P)-ONS76 Dox-induced vs control was used to create a list of genomic regions for visualization of enrichment for each group [i-SOX2-ONS76 Dox-induced vs control and i-SOX2(G76P)-ONS76 Dox-induced vs control] at those regions. Motif enrichment was measured using Homer (Heinz et al. 2010) findMotifsGenome with region sizes of 50 bp (CUT&RUN/ChIP-seq) or 200 bp (CUT&RUN/ChIP-seq/ATAC-seq).

All other data analysis and visualization were primarily conducted in the R programming environment [R Core Team (2025). R: A Language and Environment for Statistical Computing. https://www.R-project.org/] using the Tidyverse package (Wickham et al. 2019) for data munging and visualization. For ChIP-seq, ATAC-seq, and CUT&RUN, the sample correlation plots, PCA plots, and differential analysis were conducted using the DiffBind package (Ross-Innes et al. 2012) after removing ENCODE blacklist regions (The ENCODE Project Consortium 2012). Peak annotation was obtained using the Chipseeker package (Wang et al. 2022).

### Data Access

All raw and processed sequencing data generated in this study for RNA-seq, ChIP-seq, ATAC-seq, and both CUT&RUN high throughput sequencing profiles were deposited in the Gene Expression Omnibus (GSE293810, GSE293863, GSE316530, GSE316791, and GSE344670, respectively). The previously published data for FRA1 (also referred to as FOSL1) integrated in this study were obtained from the Gene Expression Omnibus (GSM1010756 and GSM2501565).

### Competing interest statement

The authors declare no competing interests.

## Acknowledgements

Heather Rizzino is thanked for editorial assistance. Funding for this project was provided by the Nebraska Department of Health and Human Services (2024-41), Team Jack Power 5 Foundation, and the Edna Ittner Pediatric Research Support Fund to AR, from the National Institute of General Medical Sciences (R00GM138920, P20GM121316) to SB, and from the National Institute of General Medical Sciences (R35GM15203) to TT. Core facilities were also supported by a NCI-Cancer Center Support Grant (5P30CA036727-39).

## Author Contributions

Conceptualization: A.R., B.D.O.G., and D.C.; Data curation and software: K.M., S.B., and T.T.; Formal analysis and visualization: B.D.O.G., K.M., E.P.M., and D.V.G.;

## Funding acquisition

A.R. and D.C.; Investigation, methodology, and validation: B.D.O.G., E.P.M., P.J.W, D.V.G., and J.P.; Resources: B.D.O.G., K.M., P.J.W., J.P., and S.B.; Supervision: A.R.; Writing – original draft: A.R. and B.D.O.G.; Writing – review & editing: All authors.

## Notes

### Competing Interest Statement

The authors have declared no competing interest.

